# Germline-targeting HIV Envelope SOSIP immunization more frequently elicits broadly-neutralizing antibody precursor responses in infant compared to juvenile rhesus macaques

**DOI:** 10.1101/2025.05.27.656273

**Authors:** Yasmine A. Issah, Xintao Hu, John Isaac, Xiaoying Shen, Dominique Davis, Gabriel Ozorowski, Leigh M. Sewall, Shiyu Zhang, Euchong Kang, Kenneth Vuong, Maria Dennis, Jui-lin Chen, John M. Ramos, Anila Yasmeen, Joshua Eudailey, Albert Cupo, Carolyn Weinbaum, Hongmei Gao, Sherry Stanfield-Oakley, Guido Ferrari, P.J. Klasse, Genevieve Fouda, Michael Hudgens, Andrew B. Ward, David C. Montefiori, Rogier W. Sanders, John P. Moore, Koen K.A. Van Rompay, Kristina De Paris, Sallie R. Permar, Ashley N. Nelson

## Abstract

A vaccine capable of inducing broadly neutralizing antibodies (bnAbs) is essential for effective prevention against HIV in children and adolescents. Germline-targeting vaccine strategies aim to stimulate bnAb precursor B cells through carefully designed immunogens, such as the stabilized SOSIP trimers, which mimic native HIV envelope (Env) proteins while presenting key neutralizing epitopes to germline B cell receptors. Given the ability of children living with HIV to develop bnAbs earlier and at a higher frequency than adults, we compared the immunogenicity of a CD4 binding site (CD4bs) bnAb germline-targeting SOSIP trimer immunization strategy in infant (n = 5) and juvenile (n = 4) rhesus macaques (RMs). Animals received 3 doses of the germline-targeting BG505 GT1.1 immunogen, followed by 3 boosts of wild-type BG505 SOSIP, each adjuvanted with the TLR7/8 agonist, 3M-052-SE. After 1.5 years, the RMs were further boosted with a mixed clade B Env trimer nanoparticle to enhance heterologous virus neutralization responses. This germline-targeting strategy induced equivalent titers of neutralizing antibodies in both groups of RMs, yet the infants exhibited a higher magnitude of vaccine-specific IgG binding. Notably, after 3 doses of BG505 GT1.1 SOSIP, infants had higher BG505 GT1.1-specific IgD^-^ B cells. Upon completion of the vaccine regimen, 4 of 5 infants developed a CD4bs bnAb precursor response detectable in serum compared to only 1 of 4 juveniles. Finally, administration of the mixed clade B nanoparticle was able to increase the breadth of antibody responses in 3 of 5 infants and 2 of 4 juveniles. These results suggest that immunization in early-life may enhance bnAb induction and highlight the potential for future pediatric HIV-1 vaccine strategies.

**Author Summary:** Vaccines that can generate antibody responses that have breadth and potency in neutralization can protect children and adolescents from acquiring multi-strain viruses like HIV. Increasingly, vaccine strategies have become better tailored to elicit these broadly neutralizing antibodies (bnAbs), such as the germline-targeting HIV envelope antigens, which have shown great promise. To determine if we could elicit protective antibody responses against HIV in early life, we immunized infant and juvenile monkeys using the same vaccine strategy.

In this study, we found that while both infants and juveniles developed similar levels of antibodies that could neutralize HIV, the infants showed stronger antibody responses in several key areas. Moreso, infants developed the type of early antibody response that researchers believe is needed to eventually evolve into broadly neutralizing antibodies.

These findings suggest that initiating HIV vaccination earlier in life may offer a better chance at generating the bnAbs needed to protect against various HIV strains. Our work highlights the potential benefits of initiating an HIV vaccine regimen in childhood.

## INTRODUCTION

Human Immunodeficiency Virus (HIV) remains a significant global health concern affecting 40 million children, adolescents, and adults worldwide. Despite efforts to reduce transmission, approximately 120,000 [83,000-170,000] infants are born with HIV each year, and 1.4 million [1.1-1.7 million] children (ages 0-14) are living with HIV as of 2023 (1–3). Moreover, the risk of HIV acquisition increases with the onset of sexual debut with approximately 360,000 new infections reported annually among adolescents aged 15-24 years worldwide (2). Ideally, developing a safe and effective HIV vaccine regimen that would be delivered from early life onwards could drastically reduce the risk of infections during adolescence and remain protective into adulthood.

A successful HIV vaccine should induce protective titers of broadly neutralizing antibodies (bnAbs) before exposure(4), given their ability to recognize and prevent diverse strains of HIV from infecting their target cells. These bnAbs mediate their activity by targeting conserved regions on the HIV envelope (Env) glycoprotein, including the CD4 binding site (CD4bs), the V2 apex, a glycan-dependent epitope in the V3 loop, epitopes at the gp120-gp41 subunit interface, and the membrane-proximal ectodomain (MPER) of gp41(5,6). These bnAbs typically emerge over time following virus acquisition, often requiring high levels of improbable somatic hypermutations (SHM) (7) combined with multiple cycles of viral escape. These factors eventually contribute to promoting increased breadth and potency of the antibody responses through affinity maturation in a subset of individuals (8–10). Interestingly, infants living with HIV can develop bnAb responses in sera faster and more efficiently compared to adults living with HIV(11–13). Further, studies have found that HIV-1 bnAbs isolated from pediatric patients have lower levels of SHM compared to those isolated from adults(13). This distinct bnAb evolution may be attributed to the fact that infant B cells display low levels of SHM that increase over time with age, reaching levels equivalent to that of adults by 6 years of age(14). Overall, this highlights a potential advantage of the early life immune landscape for eliciting bnAbs following vaccination. However, the age-related immune features that can be leveraged for optimal HIV vaccine responses have yet to be fully determined(15).

It is well established that age significantly influences immune responses to vaccines as the immune system matures over time(16). While most vaccines are administered in childhood to establish lifelong immunity, children’s immune systems differ markedly from adults. Young children typically have more naïve T cells, stronger T helper 2 (Th2) responses, higher regulatory T cell (T_reg_) levels, and reduced cellular effector functions(15). The B cell compartment in early life also has distinct features such as more naïve B cells, fewer long-lasting memory B cells, and weaker responses to both T cell-dependent and independent antigens (16). Despite these differences, emerging evidence suggests that the early life immune environment is uniquely situated for optimal vaccine responses compared to adults(17,18). For example, following immunization with an HIV envelope gp120 vaccine, children developed stronger and more durable antibody responses compared to adults(17,19). Similarly, previous studies examining T cell responses following Bacillus Calmette-Guérin (BCG) vaccination observed that infants have greater Th1 and pathogen-specific CD8^+^ T cell responses as compared to adults(20). Recently, we demonstrated that SARS-CoV-2 vaccine-elicited IgG responses in children were comparable and, in some cases, higher in magnitude than those of adults, despite children receiving a lower vaccine dose(21). Thus, HIV vaccines have the potential to be more immunogenic when administered in early life, but studies utilizing current approaches to HIV vaccination have yet to evaluate this.

One of the most promising HIV vaccine strategies employs the use of SOSIP trimers, a bio-engineered, stabilized trimeric protein designed to imitate the native-like structure of the HIV Env and has the potential to elicit bnAb responses(10,22–26). The wildtype (WT) BG505 SOSIP, based on a clade A virus isolated from an infant who developed bnAb responses(13,27) has been successful in eliciting neutralizing antibodies against both tier 1 and the more neutralization-resistant tier-2 autologous virus, as well as some low-level heterologous neutralization breadth(28). Iterative design of these trimers based on engagement of the unmutated common ancestors of bnAbs isolated from infected individuals yielded the BG505 SOSIP.v.4.1 GT1.1 (BG505 GT1.1) antigen to better engage germ-line B cell lineages and drive antibody responses specifically targeting the CD4bs and V2 apex (29). This BG505 GT1.1 SOSIP trimer has successfully elicited CD4bs-targeting bnAb precursor responses in preclinical and clinical studies, including in infant RMs(29–32).

Given the potential advantages of the early life immune environment for the development of HIV bnAbs, plasticity of the B cell responses, and the opportunity to integrate a multidose HIV vaccine strategy within the pediatric vaccine schedule, we set out to determine if there were identifiable immune advantages to implementing a HIV bnAb germline-targeting vaccine regimen during early life, including the induction of bnAb precursor responses. In this study, we compared the immunogenicity of a BG505 GT1.1 prime and WT BG505 SOSIP boost strategy in infant and juvenile rhesus macaques. We further determined whether incorporating a final boost with a heterologous multivalent HIV Env trimer nanoparticle could improve the breadth of antibody responses. Our study demonstrates that delivering a germline-targeting HIV vaccine in early life drives robust humoral immune responses and may have qualitative advantages for eliciting HIV bnAb responses compared to when these vaccine regimens are delivered later in life.

## RESULTS

### Plasma IgG and B cell responses following BG505 SOSIP immunization in infant and juvenile RMs

To evaluate age-related differences in the immunogenicity of an HIV envelope (Env) SOSIP bnAb germline-targeting immunization strategy, infant and juvenile rhesus macaques (RM) were immunized on the same schedule as outlined in **Fig 1**. Briefly, infant (median age 2 days) and juvenile (median age 2.8 years; **S1 Table**) RMs received, 3 doses at 50 μg each of the BG505 GT1.1 SOSIP (referred to as BG505 GT1.1) intramuscularly six weeks apart followed by 3 doses at 50 μg each of the wild type BG505.664 SOSIP (referred to as BG505 SOSIP) at weeks 24 (26 for the infants), 52, and 78. Immunizations were adjuvanted with 10µg of the TLR7/8 agonist, 3M052-stable emulsion (SE, 2%), a schedule previously described (32).

**Fig 1.**
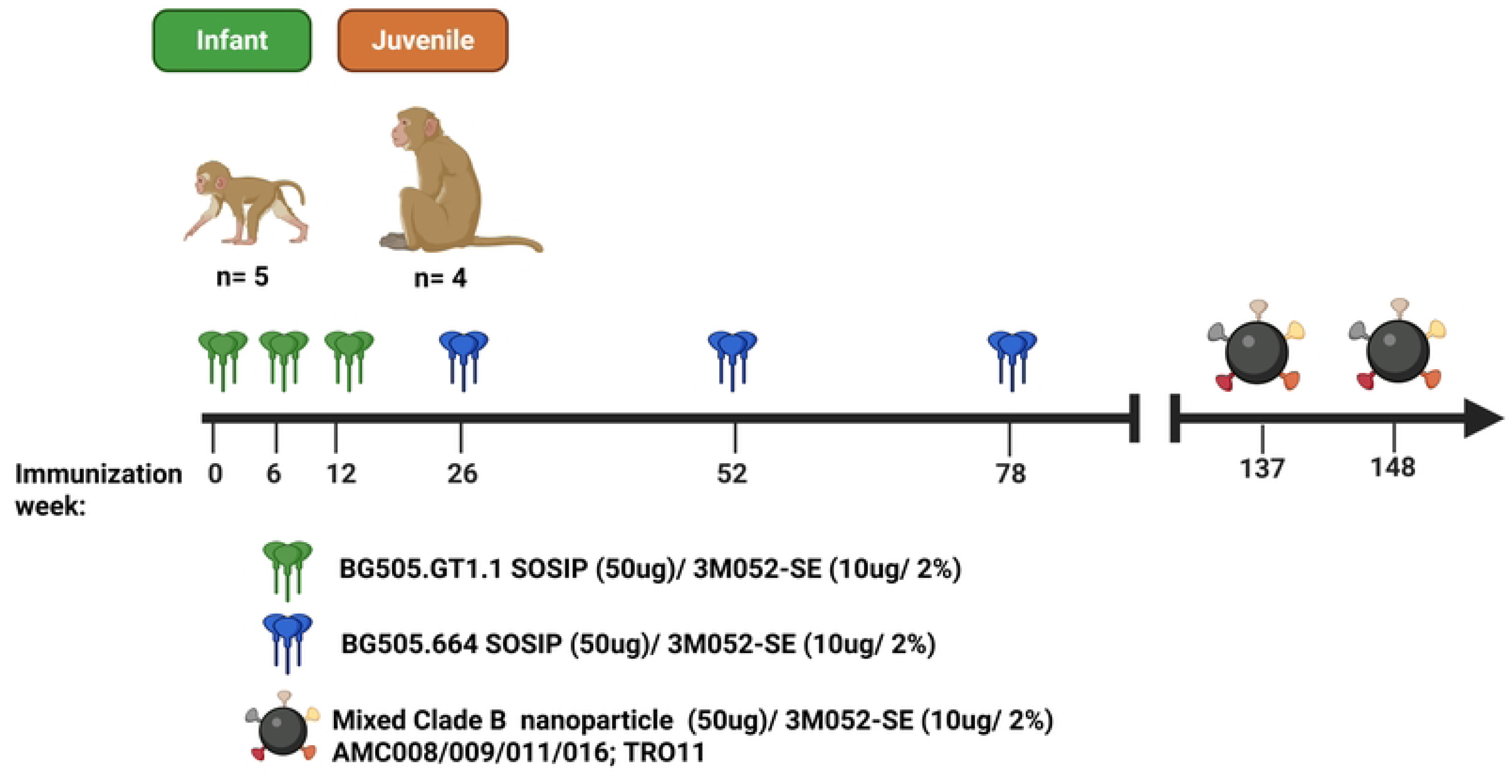
Immunization Schedule for infant (n=5; green) and juvenile (n=4; orange) rhesus macaques (RMs). Both age groups received 3 doses at 50µg each of BG505. GT1.1 SOSIP followed by 50µg of BG505.664 SOSIP. All animals received 2 doses of a mixed clade B trimer nanoparticle immunization at weeks 137 and 148. All immunizations were administered with the adjuvant 3M052-SE.

The magnitude and kinetics of vaccine-elicited IgG binding responses in plasma were assessed over the course of the study. Plasma BG505 GT1.1-specific IgG binding responses increased two weeks following each immunization in both age groups, with the peak response observed two weeks after the final BG505 GT1.1 immunization (median effective dilution at 50% response (ED_50_) at week 14: 31,051 for infants and 5,818 for juveniles; **Fig 2A**). Overall, the infant RMs consistently trended towards higher median plasma BG505 GT1.1-specific IgG binding responses compared to the juveniles throughout the course of the study (**Fig 2A**). In particular, the peak IgG binding responses trended higher in infants following each vaccine dose compared to juveniles, although levels of vaccine-specific plasma IgG antibodies declined to similar levels between doses across the age groups. Nevertheless, the median area under the curve (AUC) responses were comparable across age groups (p=0.75).

**Fig 2.**
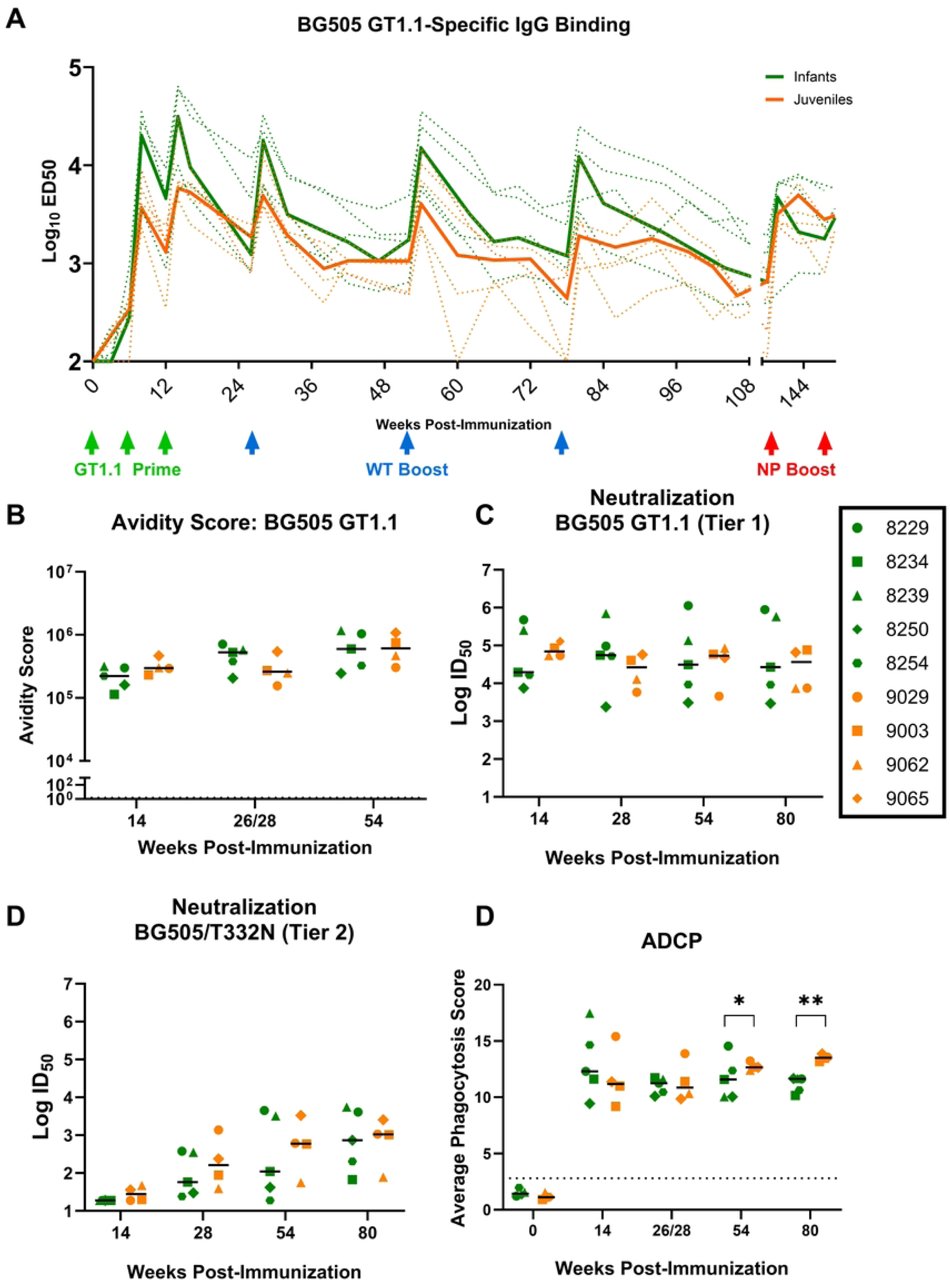
Higher plasma IgG and comparable autologous virus responses in immunized infant and juvenile rhesus macaques. (A) Plasma IgG binding responses to BG505 GT1.1 SOSIP were measured by ELISA. The bold lines represent the median; the green arrows represent GT1.1 immunization, and the blue arrows are BG505 SOSIP. (B) Plasma IgG binding avidity scores were measured by SPR against BG505 GT1.1 SOSIP protein. Serum HIV-1 neutralization assayed in TZM-bl cells against BG505 GT1.1 tier 1 virus (C) and BG505/T332N tier 2 virus (D) 2 weeks following the 3^rd^-6^th^ immunizations. (E) Plasma ADCP was measured using a flow-based assay against fluorescently tagged BG505 GT1.1-coated beads; the dotted line represents the PBS-only baseline.

The avidity of vaccine-elicited IgG binding antibodies was measured using surface plasmon resonance (SPR). IgG-purified plasma samples were screened against BG505 GT1.1 (**Figs 2B and S1**) at baseline and two weeks post-3^rd^, 4^th^, and 5^th^ immunizations (weeks 14, 26/28, and 54). At week 14, following priming with BG505 GT1.1, IgG avidity scores were similar between the two groups (median: infants-221,837 and juveniles-296,698; p=0.270). This trend continued for avidity scores at weeks 26/28 and 54 following BG505 SOSIP immunization with no significant increase with subsequent doses (median avidity score at weeks 26/28 and 54, respectively: infants-527153 juveniles-260,334 p= 0.27; infants-597,777 juveniles-610,375 p=0.90 respectively; **Fig 2B**).

### Development of serum autologous virus neutralization responses in infant and juvenile RMs following immunization with BG505 SOSIPs

Neutralizing antibody responses were first assessed in serum samples from weeks 14, 26/28, 54, and 80 (two weeks post 3^rd^-6^th^ immunizations) against the vaccine-matched virus, BG505 GT1.1 (tier 1, known to be sensitive to neutralization by early precursors of VRC01-like CD4bs bnAbs) and the parental BG505.T332N (tier 2) (**Figs 2C and 2D**). The RMs from both age groups developed high titers of neutralizing antibodies against BG505 GT1.1 by week 14 (median ID_50_ [range]: infants 19,650 [7,432-476,977]; juveniles: 69,845 [53,758-127,607]) after the initial BG505 GT1.1 prime. Following immunization with the BG505 SOSIP, infants demonstrated greater variability in BG505 GT1.1-specific neutralization titers (**Fig 2C**) compared to juveniles, yet overall median responses were similar across the two groups. Neutralizing activity against BG505.T332N was detectable in both groups beginning at week 28, suggesting that this response required at least one boost with the BG505 SOSIP. These responses continued to be modestly boosted with each dose but remained similar between the two groups. Overall, the magnitude and kinetics of plasma neutralization responses were comparable across age groups throughout the vaccine schedule (**Figs 2C and 2D**), despite infants exhibiting higher levels of antibody binding (**Fig 2A**).

### Vaccine-elicited Fc-mediated antibody responses following BG505 SOSIP immunization in infant and juvenile RMs

Although virus neutralization is the primary antibody function associated with protection from HIV-1 acquisition, antibody Fc-dependent effector functions, such as antibody-dependent cellular cytotoxicity (ADCC) and antibody-dependent cellular phagocytosis (ADCP), are increasingly recognized as important in protection against a range of viruses, including HIV-1 (33–35). Thus, we evaluated the development of vaccine-elicited antibodies capable of mediating ADCC and ADCP across the age groups (36).

Plasma was tested for ADCP responses and scored based on the proportion of BG505 GT1.1 SOSIP-coated fluorescent beads phagocytosed by THP1 cells after plasma incubation. Two weeks following the third immunization at week 14, ADCP activity was slightly higher in the infants compared to the juveniles (median phagocytosis score in infants and juveniles, respectively: 12.206 and 11.187; **Fig 2E**). However, at weeks 54 and 80, 2 weeks post-5^th^ and 6^th^ immunizations, median ADCP responses were higher in the juveniles (week 54 median: 12.47 ; week 80 median: 12.25) than infants (week 54 median: 11.58; week 80 median: 11.63), reaching statistical significance (p=0.014 and 0.0019, respectively). Furthermore, the ADCP responses remained consistent through the sixth immunization at week 80, suggesting that IgG effector function did not improve with additional doses (**Fig 2E**).

We also evaluated vaccine-elicited ADCC responses in plasma, first as NK cell activation mediated by vaccine-elicited plasma antibodies against BG505 gp120-coated targets. ADCC end-point titers were undetectable in the vaccinated juvenile RMs, yet two of five infants (8229 and 8250) developed ADCC mediating antibodies at weeks 26 and 54 (**Fig S2A**). Next, we determined the ability of vaccine-elicited antibodies to mediate ADCC against cells infected with an infectious molecular clone of BG505 (**Fig S2B**). One vaccinated infant and one juvenile RM consistently demonstrated high ADCC activity in this luciferase-based assay (**Fig S2B**).

Interestingly, one infant (8250) never developed ADCC-mediating antibodies **(Fig S2B),** and this same animal had the lowest BG505 GT1.1 neutralization titers (**Fig 2C**) and avidity score (**Fig 2B**), suggesting that this animal may not have mounted a strong functional humoral response following immunization. Across the vaccine regimen, the infants displayed more detectable, albeit transient, ADCC activity compared to the juveniles.

Glycan modifications of the Fc portion of the IgG heavy chain can modulate and potentially facilitate different Fc-dependent antibody responses like ADCC and have been linked to protection against viruses such as influenza and HIV(37–40). Thus, we evaluated glycosylation patterns of BG505 GT1.1-specific IgG across both age groups at weeks 16 and 86 of vaccination. Overall, there were no distinct differences between the plasma vaccine-elicited IgG glycosylation features in the infant and juvenile RMs except for biantennary N-glycans (G2) at weeks 84 (infants) and 86 (juveniles; p=0.04; **Fig S2C**). Further, PCA cluster analysis also showed no distinct clustering between the juveniles and infants across these two time points (**Fig S2D**). At week 16, the juveniles had a higher relative abundance of agalactosylated IgG compared to the infants (p=0.02) though this difference was not maintained by week 84/86 (**Fig S2E**). Agalactosylated IgG has been associated with pro-inflammatory disease conditions and has been found to be expanded in people living with HIV compared to healthy controls (41,42), yet its implications in vaccine-elicited Fc-mediated antibody functions remain unclear. Besides agalactosylation, there were no other distinct differences in the relative abundance of IgG glycan patterns across age groups at both weeks 16 (4 weeks post-3^rd^ immunization) and 86 (8 weeks post-3^rd^ immunization; **Fig S2E**). Thus, germline-targeting SOSIP immunization elicits IgG with similar glycan patterns across age groups, consistent with the overall lack of significant differences observed in Fc-mediated functions.

### Epitope mapping of vaccine-elicited autologous wild-type virus neutralizing antibodies following BG505 SOSIP boosting across age groups

As the BG505 GT1.1 immunogen was designed to elicit CD4bs and V2-apex-targeting bnAbs, we set out to determine the epitope specificity of vaccine-elicited neutralizing antibodies. To evaluate this, sera were screened against a panel of pseudoviruses (**S2 Table)** on the tier 2, BG505.T332N, virus backbone containing mutations corresponding to known bnAb epitopes. Epitope specificity was assigned based on a 3-fold or more decrease in sera neutralization potency against the mutant when compared to the parent (BG505.T332N) virus. Overall, we could not map the vaccine-elicited neutralizing antibodies to a single epitope on the BG505 Env in the majority of infant and juvenile RMs. However, there was one infant (8254) and one juvenile (9062) that demonstrated development of CD4bs-targeting neutralizing antibodies (**S3A Fig**).

The BG505 Env contains distinct glycan holes at residues 241/289 and 465 (43,44). These glycan holes have been found to elicit immunodominant responses against the BG505 SOSIPs, impeding the induction of CD4bs targeting antibodies and heterologous neutralization (45).

Likewise, autologous virus neutralizing antibodies elicited following BG505 SOSIP vaccination have predominantly mapped to the immunodominant C3/465 epitope, corresponding to a strain-specific glycan hole on the BG505 Env (46,47). To determine if autologous virus-neutralizing antibodies target these known glycan holes on the BG505 Env, sera were screened against pseudoviruses incorporating mutations around the C3 region glycan hole (G354E), or mutations designed to fill in glycan holes at positions C3/465 (I358T and T465N) and 241/289 (S241N.P291T; **S2 Table**). At weeks 54 and 80, three infants (8229, 8239, 8250) and two juveniles (9029 and 9065) showed tier 2 autologous virus neutralization specificity to the C3/465 epitope (**S3A and S3B Figs**). Animals 8229 and 8239 also had the highest autologous virus neutralization titers compared to other animals in the infant group at weeks 54 and 80 (**Figs 2C and 2D**). However, this trend of high autologous virus neutralization titers did not hold with the two juveniles with strong C3/465-targeted neutralization (9029 and 9065; **S3A Fig**), suggesting that high autologous virus neutralizing titers were not consistently dependent on antibody specificity for glycan holes. Further, only one infant (8234) and none of the juveniles exhibited specificity for the 241/289 glycan hole at week 54, although one infant (8254) developed this specificity at week 80 (**S3B Fig**). A decrease in specificity towards glycan holes can indicate the development of neutralizing antibodies targeting more relevant bnAb epitopes (45), although we were unable to map these specificities in our neutralization assays.

### Development of serum CD4 binding site (bs)-specific bnAb precursors and heterologous virus neutralization responses in infant and juvenile RMs

The BG505 GT1.1 immunogen was designed to engage precursors of the VRC01-class of CD4bs bnAbs (29). Therefore, we evaluated the development of these responses in serum using a previously described strategy to identify neutralization signatures of VRC01 precursors in the TZM-bl assay (48) incorporating the pseudovirus 426c.TM/GnT1 and its VRC01 knock-out mutant virus 426c.TM.D279K/GnT1-. The VRC01-precursor signature is defined by a greater than 3-fold difference in ID_50_ neutralization titers between the 426c.TM parent virus and its corresponding knock-out virus 426c.TM.D279K. By week 54 (2 weeks post 5^th^ immunization), two of five infants developed this VRC01 bnAb precursor signature, with an additional infant developing the response at week 80 (2 weeks post 6^th^ dose). However, only one of four juveniles developed this signature within the same time frame (**Fig 3A**). Interestingly, there was a trend towards higher BG505 GT1.1 neutralization ID_50_ titers in the RMs that developed the serum VRC01 precursor signature at weeks 54 and 80 (**Figs 3A and 3B**).

**Fig 3.**
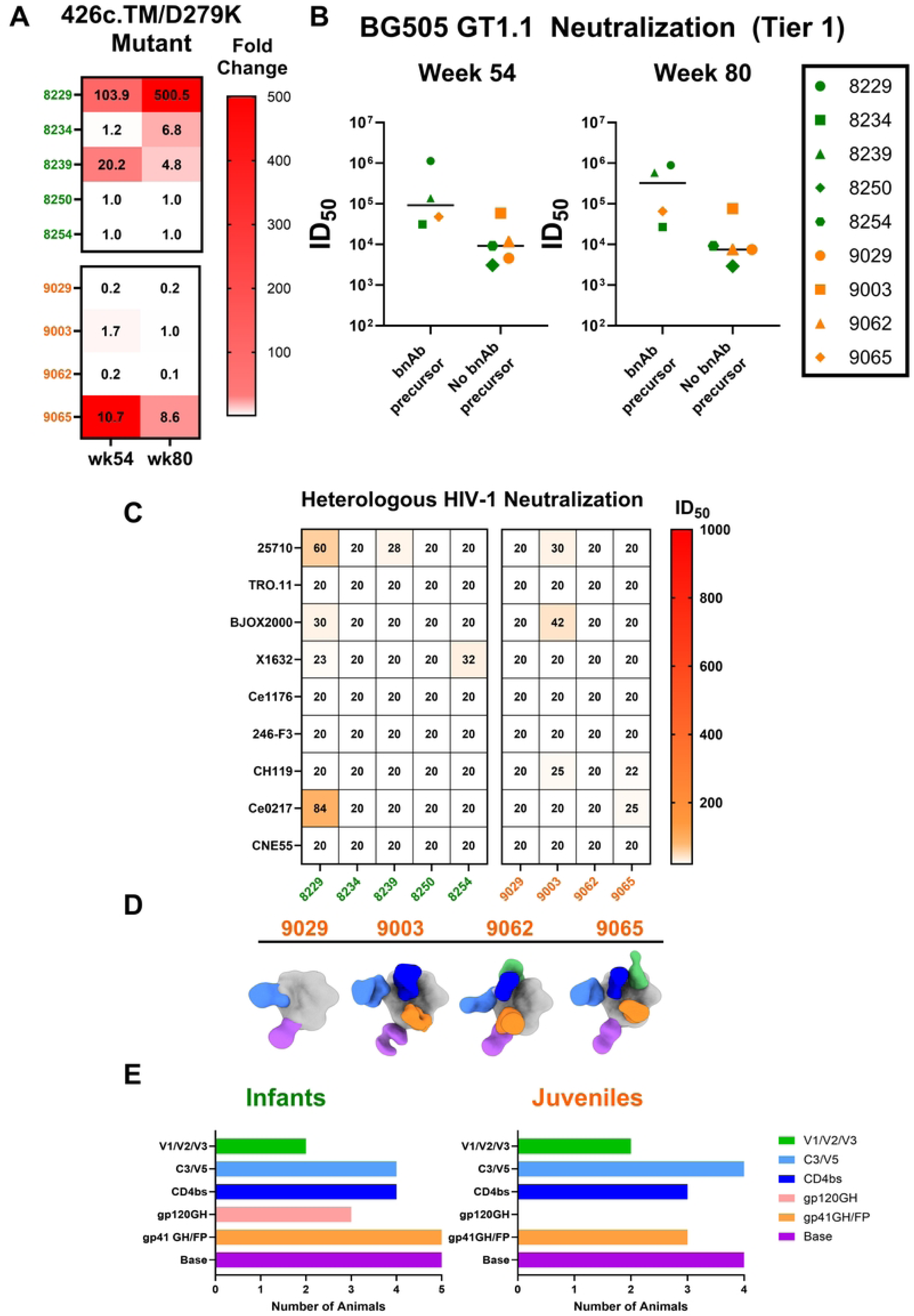
Development of CD4 binding site targeting antibodies in infant and juvenile rhesus macaques. (A) Heat map summarizing the fold change of ID50 values of sera screened against 426c.TM/GnT1-, and the CD4bs KO virus, 426c.TM.D279K/GnT1-. A VRC01-like precursor bnAb signature was assigned based on >3-fold reduction in ID50, compared to the parent virus. (B) Comparison of BG505 GT1.1 (tier 1) neutralization titers and the development of precursor bnAb response at weeks 54 and 80; infants are represented in green, and juveniles in orange. (C) Sera was screened against 9 heterologous viruses from the global HIV-1 panel for infants and juveniles at week 54 and reported as ID_50_, summarized in the heatmap. (D) nsEMPEM composite maps of plasma antibodies (colored Fabs) in complex with BG505 GT1.1 among juveniles at week 80. (E) Summary chart of nsEMPEM results, demonstrating epitopes targeted by plasma antibodies from infant and juvenile RMs; colors correspond to different epitopes targeted as outlined in the legend. GH=glycan hole (gp120:N241/N289; gp41:N611/N625), FP=fusion peptide region, CD4bs= CD4 binding site.

Sera from week 54 were also screened against a panel of HIV-1 tier 2 pseudoviruses representing global circulating isolates(49) to determine if our immunization strategy induced heterologous virus neutralization. One infant (8229) demonstrated the greatest heterologous virus neutralization breadth among both groups, neutralizing 3 out of the 9 viruses in the panel. Overall, 3 infants and 2 juveniles developed antibodies that neutralized at least 1 heterologous virus in the panel, nevertheless, these few cases of heterologous tier 2 virus neutralization were of low potency as ID_50_ titers were <100 (**Fig 3C**).

### Negative stain EM epitope mapping of polyclonal antibody responses

Infant antibody responses following natural HIV-1 infections are generally more polyclonal in nature (50). To explore the epitope-specificity of the vaccine-elicited antibodies, we used negative stain electron microscopy polyclonal epitope mapping (nsEMPEM) to visualize plasma IgG binding to potential neutralization epitopes on BG505 GT1.1 Env at week 80 (2 weeks post 6^th^ immunization) (**Fig 3D**). A majority of animals across age groups had detectable polyclonal antibodies to 4 or more distinct epitopes on the HIV Env trimer, with the most common target being the CD4bs, gp41-base, C3/V5, and gp41 fusion peptide region (**Figs 3D and 3E**). Some of the infant RMs also showed polyclonal antibody responses against the gp120 N241/N289 glycan hole (8234, 8239, and 8250), while none of the juveniles developed responses against that strain-specific epitope (51). The nsEMPEM results support the CD4bs-specific IgG in RMs that developed the CD4bs bnAb precursor neutralization signature, which included the RMs that displayed the highest diversity in epitope binding in the nsEMPEM (8229, 8234, 8239, and 9065) (**Fig 3E**), as previously reported (32). Moreover, the infant RMs developed antibodies targeting a greater variety of epitopes on the BG505 GT1.1 Env compared to juvenile RMs (**Fig 3E**), suggesting that HIV vaccination also elicits a more polyclonal antibody response when administered in early life.

### Early kinetics of vaccine-elicited peripheral B cell responses

We next assessed the circulating B cell populations at week 14, 2 weeks post-3^rd^ immunization with BG505 GT1.1, when the biological age differences between the groups should be the most pronounced (**S4A Fig**). Despite no significant differences in the frequency of circulating CD20^+^ B cells (median: infants 14.20% and juveniles 14.90%; p=0.90) across age groups (**S4B Fig**), infants had trend towards higher frequency of circulating plasmablasts (defined as CD3^-^CD14^-^ CD16^-^CD20^-^CD138^+^) compared to juveniles (median: infants-2.54% and juveniles-1.00%; **S4C Fig**). Other B cell populations were comparable between the age groups, yet non-class switched IgD^+^ (median: infants-89.00% juveniles-81.00%; p=0.39) and CD21^+^CD27^+^memory (median: infants-0.60% juveniles-0.19%; p=0.04) B cell populations displayed a higher frequency in vaccinated infants versus juveniles (**S4D and S4H Figs**). When assessing BG505 GT1.1-specific B cell responses, infant and juvenile RMs had comparable proportions of antigen-specific total and IgD^+^ B cell populations (p=1.00; **Figs 4A and 4B**). However, the proportion of circulating BG505 GT1.1 antigen-specific IgD^-^ CD20^+^ B cell subset was significantly higher in the vaccinated infants compared to that of juveniles (median: infants-1.29 juveniles-0.44; p=0.02; **Fig 4C**).

**Fig 4.**
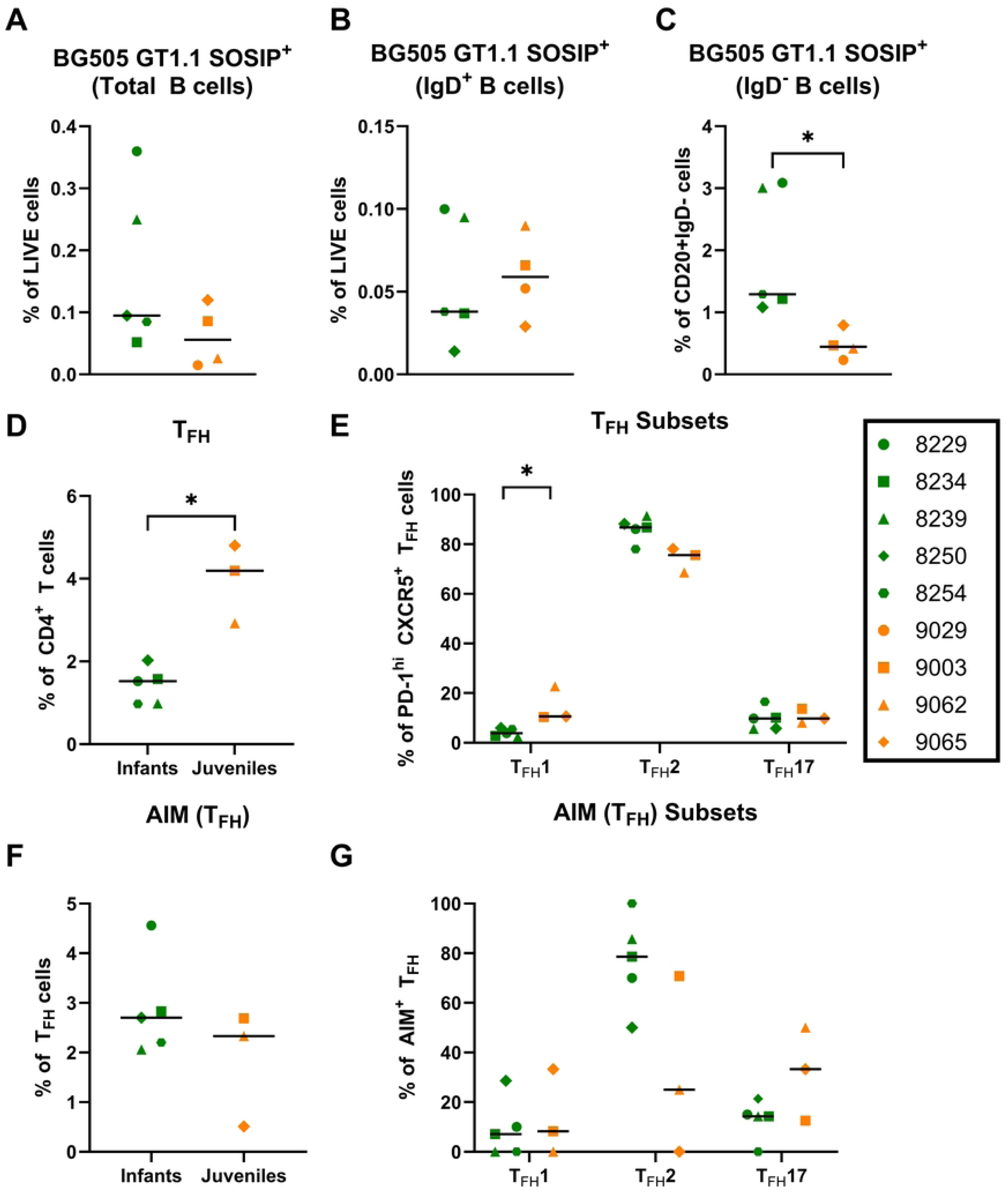
Distinct T_FH_ cell responses in infant and juvenile rhesus macaques following BG505 GT1.1 SOSIP immunizations. PMBCs were isolated from the blood at week 14 and measured for BG505 GT1.1-specific B cells gated from (A) the total CD20+ B cells population, (B) the IgD^+^ population, and (C) the IgD^-^ population. Mononuclear lymph node cells from week 54 measured for (D)Proportion of T_FH_ cells defined as a percentage of Live, CD4+ cells; T_FH_ cells: CD20^-^CD3^+^CD4^+^PD-1^+^CXCR5^+^CD134^+^CD137^+^ (E) T_FH_ subsets defined by their surface marker expression (T_FH_ 1:CD183(CXCR3)^+^CD196(CCR6)^-^; T_FH_ 2: CD183(CXCR3)^-^ CD196(CCR6)^-^; T_FH_ 17: CD183(CXCR3)^-^CD196(CCR6)^+^). (F) Frequency of antigen-specific T_FH_ cells evaluated via AIM assay; gated from Live, CD4^+^ cells. (G) Frequency of antigen-specific T_FH_ subsets. Lymph node PBMCs were used from both groups at week 54.

### Vaccine-elicited peripheral and lymph node T cell responses

T follicular helper cell (T_FH_) responses are associated with the development of neutralization breadth in children and adults living with HIV (52,53). Therefore, we measured T_FH_ cell responses in lymph nodes and peripheral blood by flow cytometry (**S3 Table**) at week 54 (2 weeks post the 5^th^ immunization), when CD4bs-specific bnAb precursors were first observed in both age groups. T_FH_ cells were initially identified as CD20^-^CD3^+^CD4^+^CXCR5^+^PD-1^hi^, and then further examined by subsets defined as: T_FH_1: CCR6^-^CXCR3^+^, T_FH_2: CCR6^-^CXCR3^-^, T_FH_17: CCR6^+^CXCR3^-^. Overall, the proportions of T_FH_ cells in the lymph nodes were higher in juvenile RMs compared to the infants (median: infants 1.52% and juveniles 4.19%; P=0.04; **Fig 4D**). Within this cell population, juvenile RMs had a higher proportion of T_FH_ type 1 cells (median: infants 3.87% and juveniles 10.60%; P=0.04), while infants trended towards higher T_FH_ type 2 cells (median: infants 86.80% and juveniles 75.60%; p=0.07), and no difference was observed in T_FH_17 proportions between the groups (**Fig 4E**). These findings are consistent with previous studies, where infant RMs displayed a higher proportion of T_FH_ type 2 responses while adult RMs displayed higher T_FH_ type 1 responses following SHIV infection(18). The infant and juvenile RMs displayed a comparable proportion of antigen-specific T_FH_ cells defined by additional markers, OX40^+^CD137^+^ (median: infants 2.70% and juveniles 2.33%; p=1.00; **Fig 4F**), and measured by the activation-induced marker (AIM) assay. Interestingly, the infants maintained a higher proportion of T_FH_ type 2 cells compared to the juvenile RMs within this antigen-specific population (median: infants 78.60% and juveniles 25.00%; p=0.14; **Fig 4G**).

We also characterized the vaccine-elicited CD4^+^ and CD8^+^ T cell responses against pooled BG505 Env peptides (**S3 Table**) at late time points (weeks 92-94) in the vaccine regimen. At this time, an adequate amount of PBMCs were available in both groups, and the immaturity of the cellular immune system in infants is less pronounced, thereby allowing us to assess the durability of this response. The infant RMs demonstrated a higher proportion of Env-specific CD4^+^ T cells than the juveniles (median: infants 63.20% and juveniles 49.15%; p=0.07), and accordingly, juveniles trended higher for the proportion of Env-specific CD8^+^ T cells (median: infants 27.60% and juveniles 38.15%; p=0.04; **S5A and S5C Figs**). Yet, there were no significant functional differences in vaccine-elicited CD4^+^ and CD8^+^ T cells based on cytokine markers, except for CD4^+^ TNFα^+^ in which the infants had a higher proportion (median: infants 0.08% and juveniles 0.00% ; p=0.03; **S5B and S5D Figs**).

### Improved heterologous neutralization responses following late boost with a mosaic HIV Env nanoparticle

To improve the breadth of vaccine-elicited neutralizing antibody responses, the infant and juvenile RMs received a ferritin nanoparticle (NP) displaying five trimeric clade B HIV-1 Envs— AMC008, AMC009, AMC011, AMC016, and TRO11, at week 137 after the initial vaccine (**Fig 1**). This pentavalent NP was previously shown to boost both autologous and heterologous virus-binding antibodies (25).

To first assess the immunogenicity of the NP boost, we tested the binding of plasma IgG against the trimeric clade B antigens using a multiplexed bead-based antibody binding assay at weeks 137 (pre-NP boost), 139, and 150 (two weeks following the 1^st^ and 2^nd^ NP boost) (**Figs 5A and S6**). The IgG binding responses to SOSIP trimers increased, as expected, following the first NP boost, though binding levels did not increase after the 2^nd^ boost at week 150 (**S6A Figure**). A similar trend was observed for BG505 GT1.1-specific binding responses (**Fig 2A**). The infant and juvenile RMs had comparable plasma IgG binding responses across the individual NP Env trimer antigens (**Figs 5A and S6**).

**Fig 5.**
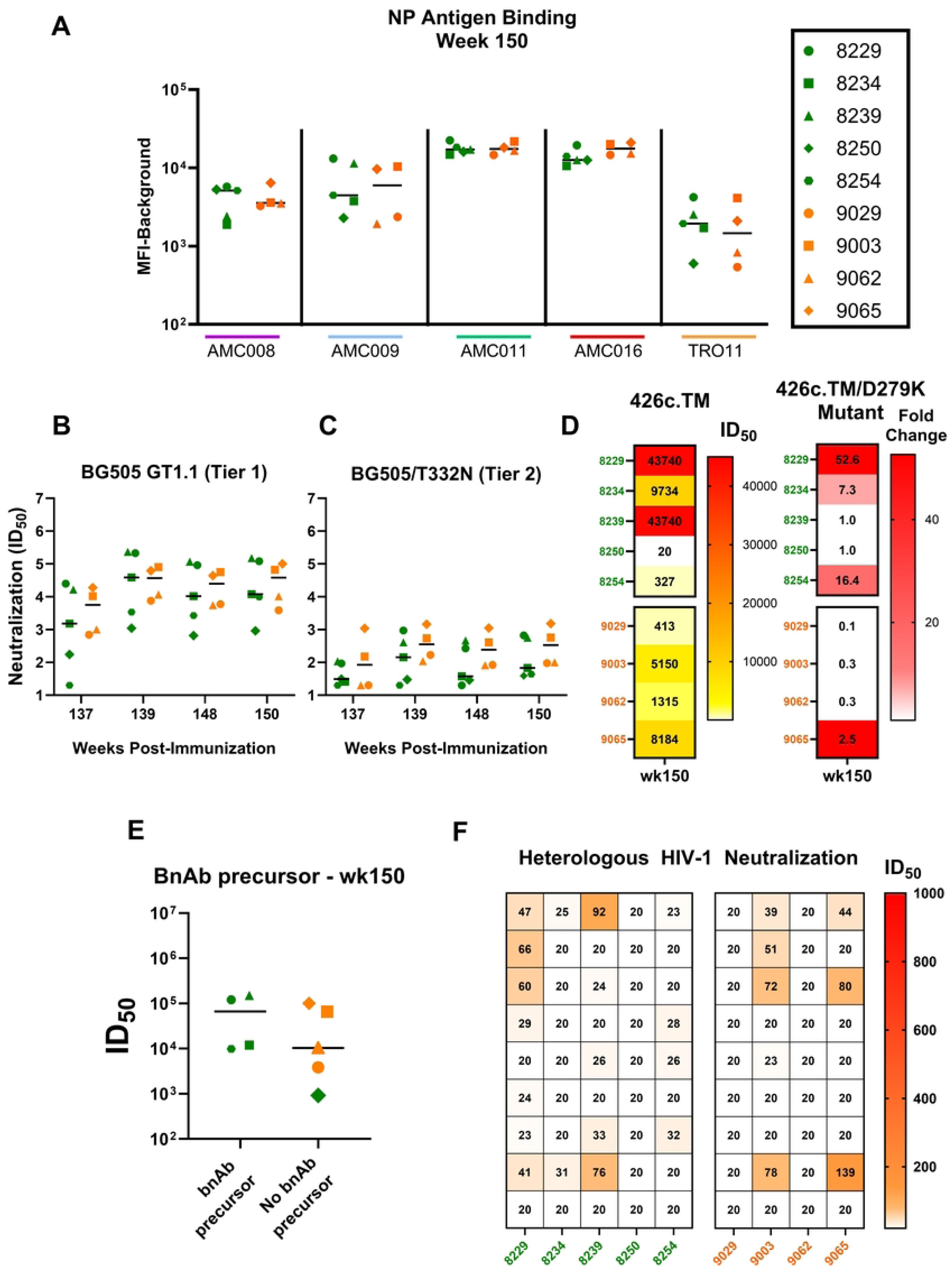
Improved breadth of vaccine-elicited antibody responses post-NP boost. (A) Plasma antibody binding tested against clade B nanoparticle vaccine antigens AMC008, AMC009, AMC011, AMC016, and TRO11 at week 150; binding responses reported in MFI (background subtracted) (B) Serum neutralization tested against tier 1 BG505 GT1.1 (B), and tier 2 BG505.T332N (C) from weeks 137-150. Neutralization titers are reported as ID_50_. (D) Development of CD4bs bnAb precursors were evaluated by screening serum against 426c.TM (parent) and 426c.TM.D279K (mutant) at week 150; titers were reported in ID50 (left panel) and fold reduction between WT and Mutant (right panel). (E) Prediction of the development of precursor bnAb response at week 150 based on the magnitude of BG505 GT1.1 (tier 1) neutralization titers; infants are in green and juveniles in orange. (F) Heatmap summarizing serum neutralization titers against a panel of 9 heterologous viruses for infants and juveniles at week 150.

Neutralization responses against the autologous, BG505 GT1.1 and BG505/T332N viruses (**Figs 5B and 5C**) were boosted by the multi-Env NP immunization yet did not exceed the magnitude achieved after the initial SOSIP vaccine doses (**Figs 2C and 2D**). Moreover, there were no differences in neutralization titers observed between the infant and juvenile RMs (**Figs 5B and 5C**). Interestingly, after the NP boost, an additional infant (8254) demonstrated a >3-fold reduction in sera neutralization titers when comparing between the 426c.TM parent virus, and its corresponding mutant, 426c.TM.D279K. Thus, a total of 4 out of the 5 infants developed the VRC01-like precursor signature in sera over the course of the study (**Fig 5D**). Unlike the infant RMs, no additional juvenile RMs developed the plasma CD4bs bnAb precursor signature after the multi-Env NP boost. Furthermore, the juvenile RM that had previously achieved this response by week 80 (9065; **Fig 5A**) did not maintain this signature at week 150. Interestingly, there was no difference in the autologous tier 1 BG505 GT1.1 neutralization response between those RMs that developed the CD4bs bnAb precursor response (**Fig 5E**), as previously observed with the earlier time points (**Fig 3B**). Finally, following the second NP boost, 4 of 5 infants had some moderate improvement of breadth in their heterologous virus neutralization responses, while only 2 juveniles showed improvement in breadth (**Fig 5F**).

## DISCUSSION

Developing vaccines for children and adolescents that induce life-long immunity leverages a critical window for immunization, potentially leading to long-term protection throughout adulthood and reduced transmission within communities. A powerful example is the rubella vaccine(54), which when administered in early childhood, provides lifelong immunity and prevents infection during pregnancy, thereby eliminating severe fetal consequences. An ideal HIV vaccine would similarly elicit immunity and provide protection prior to sexual debut, necessitating administration in childhood. In this study, we compared the immunogenicity profiles of an HIV vaccine designed to elicit difficult-to-induce bnAb responses. We evaluated the combined effects of a bnAb germline-targeting BG505 GT1.1 SOSIP, a wild-type BG505 SOSIP, and a mixed Clade B Env trimer nanoparticle in infant and juvenile rhesus macaques, all of which received the same multidose immunization regimen over 150 weeks. Our findings demonstrate that early-life immunization is highly immunogenic and may offer advantages for bnAb induction compared to starting this vaccine regimen later in life. Notably, BG505 GT1.1 prime/BG505 SOSIP boost strategies successfully elicited high titers of serum neutralizing antibodies that exhibited a neutralization phenotype indicative of early precursors of VRC01-like CD4bs bnAbs.

We established that both the infant and juvenile RMs mounted a robust binding antibody response to the BG505 GT1.1 prime and BG505 SOSIP boost. Though the infants consistently demonstrated a higher magnitude of plasma IgG antibody binding to BG505 GT1.1 SOSIP antigen, the autologous neutralization titers for both the germline targeting and the parental BG505 viruses were comparable across age groups, suggesting that this immunization regimen elicits similar epitope-specific and functional antibody responses. This finding is in line with earlier human infant studies, which established that an infant HIV-1 gp120 vaccination was able to elicit durable and high-magnitude IgG responses against specific epitopes on the HIV Env (55). Similarly, in the context of SARS-CoV-2 vaccination, infants generate similar to or even higher magnitude binding and neutralizing IgG antibodies to a lower dose of vaccine compared to older children (56–58). Moreover, these antibody responses are quite durable compared to adults (59). Infants infected with SARS-CoV-2 have also been shown to have higher levels of ADCC compared to their infected mothers (60). One possibility for the mechanism behind higher magnitude plasma IgG responses elicited in infants compared to older ages is that infant B cell responses are skewed toward antibody-producing plasmablasts compared to memory B cells (61,62). In this study, we found no significant differences in most circulating B cell populations, however, plasmablasts and antigen-specific BG505 GT1.1 SOSIP B cells trended higher for the vaccinated infants compared to those of juveniles (**S2 Fig**).

Plasma IgG epitope mapping studies in infants living with HIV have identified that infants frequently generate polyclonal HIV-neutralizing responses targeting epitopes, including the V1V2, V3-glycan N332, and the CD4bs (63). Indeed, we recently demonstrated that the germline-targeting BG505 GT1.1 SOSIP strategy was able to elicit antibodies that targeted the CD4bs regions in infant RMs (32). Moreover, we further validated our epitope mapping using nsEMPEM, showing that this germline-targeting strategy was able to elicit polyclonal responses that targeted key conserved sites at the CD4bs and the Apex (V1/V2/V3 combined) in both age groups. It has been established that infants living with HIV can develop polyclonal, cross-clade nAb responses by 20 months of age because the HIV Env can activate multiple B cell lineages (12). The only difference in the epitope-specificity elicited in infants compared to juvenile RMs detected by EMPEM was an additional antibody epitope specificity to the N241/N289 strain-specific glycan hole compared to the juveniles. Overall, the juvenile RMs in this study mounted a similar polyclonal response to epitopes on the BG505 GT1.1 Env trimer to that of infants, though the epitope-specific responses were more often universal in infants. This further supports previous observations suggesting infants living with HIV can mount HIV-1 specific neutralizing antibodies targeting the same key bnAb epitopes as adults.

The ability to engage bnAb precursor B cells and guide their evolution through shaping and refining HIV vaccine antigens is crucial for consistently eliciting bnAb responses in plasma (64). An important difference in vaccine responses between age groups was that this germline-targeting immunization strategy successfully elicited precursors of CD4bs-specific VRC01-like bnAbs in the plasma of four out of five infant RMs, whereas only one out of four juvenile RMs developed this response. Indeed, Caniels et al. recently reported VRC01-like bnAb precursor development in only 50% (n=3 of 6) of adult RMs following five immunizations with GT1.1 and WT BG505 SOSIP (29). Given the differences in vaccine dose and adjuvant between the Caniels et al. study and our own, these results reveal important considerations for the administration of an HIV vaccine across different age groups. Depending on age, the modification of vaccine antigen dose, adjuvant selection, and timing between boosts can be tailored to target bnAb precursor activation with higher consistency. Nonetheless, our observations support the strategy of initiating germline-targeting vaccination early in life to select and expand precursor B cells capable of ultimately producing bnAbs after subsequent immunizations.

In the context of viral infection, adult and infant rhesus macaques respond to SHIV infection with similar kinetics in the development of plasma-neutralizing antibodies yet demonstrate differences in T follicular (T_FH_) and germinal center B cell frequencies in lymphoid tissues (18). Within 12 weeks post-infection, infants generated higher proportions of T_FH_ and germinal center B cells in the lymph nodes compared to adults (18). Accordingly, the juvenile RMs displayed higher frequencies of total lymph node T_FH_ cells than infants after SOSIP boost vaccination, indicating less robust germinal center responses following vaccination in infants. The early life immune system is characterized by Th2-skewed and modest effector T-cell responses. Accordingly in our study, infants demonstrated higher Th2 and lower Th1-type vaccine-elicited T_FH_ cell responses. Future work should explore adjuvants that could overcome the limited T cell response, peripherally and in the germinal centers, to promote rare vaccine-elicited B cell evolution in infants. Additional strategies to optimize dosing in pediatric populations for optimal cellular and humoral immunity should be explored as well. Given the distinct nature of immune environments during development in childhood, understanding the ontogeny of immune responses to germline-targeting vaccines will be critical for refining vaccination protocols that will leverage the uniqueness of the immune responses at specific age groups to induce bnAbs effectively across the lifespan.

Studies of bnAbs isolated from infants living with HIV revealed that one V3 glycan-specific bnAb, BF520.1, displayed characteristics of cross-clade, heterologous neutralization within 6 months of infection and remarkably low levels of SHM (65). This contrasts with previously isolated bnAbs from adults, where those same responses take many years to develop. Recently, SHIV infection models in infant rhesus macaques have also established that early life immunity results in heterologous HIV-1 neutralization responses (66). Additionally, there have been other reported instances of infants having higher responses to heterologous viruses compared to adults, such as with influenza (67). We included a mixed clade B nanoparticle boost at the end of the nearly 2-year vaccine schedule that simulated pre-adolescent vaccination in the younger RM group, seeking to improve the breadth of cross-clade neutralization responses. We found that the two boosts with the mixed clade B nanoparticle maintained the VRC01-precursor bnAbs in three infants and boosted this response to be detectable again after 2 weeks in one animal at week 150. Furthermore, the nanoparticle boosts moderately improved the breadth of heterologous neutralization, especially in the infant RMs. Following the first nanoparticle boost, the plasma antibodies displayed comparable binding to the nanoparticle antigens and were able to elicit high neutralization titers against tier 1 BG505 GT1.1 virus in both age groups. However, the second boost did not further increase antibody binding and neutralization titers. Thus, the inclusion of the mixed clade B nanoparticle vaccine boosts to improve the breadth of germline-targeting HIV vaccine strategies could potentially rely on a single dose to improve autologous and heterologous neutralization.

While our immunization strategy points to advantages in implementing an HIV-1 vaccine in early life, further assessment of how early antibody responses observed in this study translate into durable immunity against natural infection or if additional immunogens are required to generate and maintain protective immunity is needed. The relatively small sample size limits the generalizability of the findings. Future studies should include more animals to confirm these results and assess the impact of dose and adjuvant choice. The age-dependent differences in immune responses observed in this study have important implications for HIV vaccine development and the appropriate target populations. Moreover, the BG505 GT1.1 SOSIP vaccine strategy is designed to target both the CD4bs and V2-apex bnAb lineages. Antibody responses against both epitopes have been demonstrated to be some of the broadest and potently neutralizing against HIV (68,69). Future immunogen design efforts should continue incorporating modifications that engage multiple bnAb lineages and assess whether concurrent or serial introduction of these multi-epitope germline-targeting vaccines should be designed. While our studies were limited to the analysis of serum antibodies, future studies should examine the differences in the antigen-specific B cell repertoire elicited across ages to determine whether distinct B cell receptor mutation patterns emerge in the repertoire that are associated with the timing of vaccine initiation. Furthermore, this may allow us to explore the mechanisms behind the rapid and frequent induction of bnAb responses in early life, which could be better exploited through bnAb germline-target vaccination.

In conclusion, this study points to the potentially advantageous strategy of early life immunization for achieving vaccine-elicited protective HIV immunity across the age span. Initiating the vaccine in the juvenile age group, akin to the current adolescent vaccine schedule, was not as successful in initiating a critical bnAb lineage as initiating the vaccine in early life. These findings support the continued exploration of early-life vaccination strategies, with a focus on optimizing immune responses through tailored vaccine regimens for different immune development stages. Furthermore, these studies offer valuable insights for the design of clinical trials aimed at optimizing germline-targeting HIV immunization strategies in human infants and adolescents, aiming to achieve protective immunity prior to the highest risk periods for sexual exposure to the virus.

## MATERIALS AND METHODS

### Animal Information and Ethics Statement

Type D retrovirus-, SIV- and STLV-1 free infant (n=5) and juvenile (n=4) Indian origin rhesus macaques (RM) (Macaca mulatta) were maintained in the colony of the California National Primate Research Center (CNPRC, Davis, CA). Animals were maintained in accordance with the American Association for Accreditation of Laboratory Animal Care standards and The Guide for the Care and Use of Laboratory Animals. All protocols were reviewed and approved by the University of California at Davis Institutional Animal Care and Use Committee (IACUC) prior to the initiation of the study.

### Immunization Schedule and Sample Processing

Animals were immunized via intramuscular injection three times in 6-week intervals with 50µg of BG505 SOSIP.v4.1-GT1.1 (abbreviated to BG505 GT1.1) followed by boosts at weeks 24, 52, and 78 with BG505.664 SOSIP (50µg)/ 3M052-SE (10µg/2%). Lastly, at weeks 137 and 148 all animals were given a mixed clade B nanoparticle consisting of the AMC008, AMC009, AMC011, AMC016, and TRO11 HIV Env antigens. All immunogens were delivered with 10µg 3M-052 adjuvant in a 2% squalene emulsion (3M-052 SE; provided by AAHI and 3M). For immunizations and sample collections, animals were sedated with ketamine HCl (Parke-Davis) injected at 10mg/kg body weight. Blood (either without an anti-coagulant or with EDTA as an anti-coagulant was collected via peripheral venipuncture. Peripheral lymph node biopsies (of the axillary or the inguinal lymph nodes) were collected during sedation via surgery with additional local anesthesia and followed by post-surgery analgesia according to established CNPRC Standard Operating procedures.

Serum or plasma was separated from whole blood by centrifugation at 900xg for 10 minutes at 20°C. After centrifuging, plasma was collected to within 10 mm of the buffy coat and centrifuged once more at 2000xg for 15 minutes at 20oC to pellet any extracellular debris, and the plasma supernatant was frozen in multiple aliquots at -80°C. The rest of the EDTA-coagulated blood was used to isolate PBMCs by density gradient centrifugation using Ficoll®-Plaque (Sigma). Cells were then washed, given ACK lysis buffer to remove any remaining red blood cells, and cryopreserved for future use.

### Enzyme-linked immunosorbent assay (ELISA)

His-Tagged BG505 GT1.1 SOSIP (1.5-2ug/mL per well) were captured on a Nickel-coated 96 well plates (Qiagen) overnight in 1X TBS at 4°C. Plates were washed with 1x TBS and blocked for 0.5-1 hour in blocking buffer (1xTBS, 2% skim milk (VWR), and 20% Goat Serum (Sigma)). Plasma was prepared 1:100 and serially diluted 3-fold in diluent, along with recombinant mAbs B12R1 (rhesusized CD4 binding site mAb; 1 µg/mL) PGT151 (1 µg/mL) and 17b (1 µg/mL) used as controls. Plasma and mAb dilutions were added to the plates and incubated for an hour at room temperature. Following another wash, the secondary polyclonal goat anti-monkey IgG horseradish peroxidase (HRP)-conjugate (abcam) was prepared 1:10,000 and added to each well, and incubated for an hour at room temperature. The plate was washed and 100 µl TMB substrate (SureBlue; VWR) was added and incubated for 2.5 minutes at room temperature. Finally, 100 µl of stop solution is added and plates were read at an absorbance of 450 nm on the Biotek Synergy Plate Reader (Agilent).

### Virus Neutralization Assay

Antibody-mediated virus neutralization was carried out in a validated luciferase reporter gene-assay in TZM-bl cells (70). A pre-titrated dose of virus was mixed with serial 5-fold dilutions of serum samples or mAbs in duplicate for a total of 8 dilutions and incubated at 37°C for 1hr before adding TZM-bl cells. Neutralization was measured two days later as a function of reductions in luminescence. Neutralization titers are the 50% and 80% inhibitory dilution (ID_50_/ID_80_) of serum and inhibitory concentration (IC_50_/IC_80_ in µg/mL) of mAb. Assays were performed with pseudoviruses containing Env corresponding to either the BG505 GT1.1 priming immunogen (tier 1 phenotype) or the BG505/T332N boosting immunogen (tier 2 phenotype). Neutralization signatures of early precursors of VRC01-like antibodies were identified by differential neutralization of the 426c.N276D.N460D.N463D virus and corresponding VRC01 knockout mutant virus (426c.N276D.N460D.N463D.D279K), both produced in GnT1-cells to enrich for Man5 glycoforms of N-linked glycan that otherwise are processed into large complex-type glycan (48). Mapping of epitope specificity and glycan hole targeting was performed against a panel of viruses listed in **S2 Table** (23). Heterologous virus neutralization was measured by testing serum samples against 9 viruses from the global panel of HIV (49).

### Antibody-dependent Cellular Cytotoxicity (ADCC) Assays

ADCC assays were conducted as previously described (36). Briefly, ADCC activity of plasma antibodies was measured using a flow-based GranToxiLux (GTL) Assay where the cut-off for positivity in the GTL assay was >8% of Granzyme B activity. Plasma samples were serially diluted, starting at 1:100. Additionally, ADCC activity in plasma was assessed using a Luciferase-based assay where CEM.NKR_CCR5_ cells were infected with a BG505 Infectious molecular clones (IMC). For the analysis, the background was subtracted based on background detected pre-immunized samples. After removing the background, the results were considered positive if the % specific killing was above 10%. ADCC was reported in antibody endpoint titers, which were determined by interpolating the last positive dilution of the plasma (>10% Specific Killing).

### Antibody-dependent cell phagocytosis (ADCP) assay

Biotinylated BG505.SOSIP v4.1 GT1.1 antigen were conjugated to fluorescent NeutrAvidin beads (Invitrogen, Cat# F8776). Positive control mAbs were diluted at 50 ug/ml and plasma samples at a 1:50 dilution were added with antigen-conjugated beads and then incubated plate for 2 hours at 37°C with 5% CO_2_ in a 96-well plate to allow the formation of immune complexes. After incubation, a human leukemia monocytic cell line, THP-1 cells (ATCC, Cat# TIB-202) were added and centrifuged at 1,200 g for 1 hour at 4°C. To allow phagocytosis to happen, the plate was incubated for no more than 1 hour at 37°C with 5% CO_2_ following centrifugation. THP-1 cells were then fixed with 4% paraformaldehyde (Sigma, Cat# J61899-AP), and the populations of the fluorescence cells were analyzed by flow cytometry (LSRFortessa; BD). A negative control, containing a 0.1% mixture of phosphate-buffered saline and bovine serum albumin solution, was used to normalize the background phagocytosis activity. The phagocytosis scores were formulated by multiplying the mean fluorescence intensity (MFI) and frequency of positive cells and divided by the negative control’s MFI and frequency of positive cells. The samples were tested in two replicate assay and the average phagocytosis score were reported.

### Negative Stain EMPEM

Serum processing and sample preparation to obtain polyclonal fabs for electron microscopy were previously described in (PMID: 35507649). Briefly, IgG was isolated from 0.5 mL of sera taken at week 80 from BG505 GT1.1 immunized infant (n=5) or juvenile (n=4) RMs using Protein G (Cytiva). Papain (Sigma Aldrich) was used to digest IgG to fabs. 15 µg of BG505 SOSIP or GT1.1 SOSIP trimer was mixed with 1 mg of Fab mixture (containing Fc and residual papain) for an overnight incubation and the complex was then purified using a Superdex 200 Increase 10/300 GL gel filtration column (Cytiva). Purified complexes were concentrated and diluted to a final concentration of 0.03 mg/mL. The diluted samples were deposited on glow-discharged carbon-coated copper mesh grids, followed by staining with 2% (w/v) uranyl formate. Electron microscopy images were collected on an FEI Tecnai Spirit T12 equipped with an FEI Eagle 4k x 4k CCD camera (120 keV, 2.06 Å/pixel) and processed using Relion 3.0 (71) following the standard 2D and 3D classification procedures. UCSF Chimera (72) was used to generate the composite maps, and representative maps of unique epitopes identified across the study have been deposited to the Electron Microscopy Data Bank.

### B-cell Phenotyping

Cryopreserved PBMCs were thawed and resuspended in media (RPMI, 10% FBS; cells were then counted and assessed for viability. Afterward, cells were spun down and resuspended into 1xPBS + 1%BSA (VWR). A total of 1×10^6^ cells were stained with a viability dye for 30 minutes at room temperature and then washed with 1xPBS. The cells were then stained with an antibody cocktail (see **S3 Table** for a list of antibodies) for 20-30 minutes at room temperature. To stain for HIV-antigen specific cells, a tetramer was made from biotinylated BG505 GT1.1 conjugated to a streptavidin-tagged fluorophore and included in the antibody panel(73). Cells were then washed with 1x PBS and then fixed with 10% formalin in 1xPBS. Compensation was done with beads (Thermo Fisher) that were incubated with each fluorophore; Fluorescent Minus One (FMO) controls were made to aid in the gating strategy. Utilizing the gating strategy outline in **S4A Fig**, this panel was used to distinguish populations of total B cells (CD3^-^CD14^-^CD16^-^ CD20^+^), plasmablasts (CD3^-^CD14^-^CD16^-^CD20^-^CD138^+^), class-switched (CD3^-^CD14^-^CD16^-^ CD20^+^IgD^-^), and non-class-switched (CD3^-^CD14^-^CD16^-^CD20^+^IgD^+^) B cells(18,32). From the IgD^+^, subsets were divided into naïve (IgD^+^CD21^+^CD27^-^) and unswitched memory (IgD^+^ CD21^-^ CD27^+^), while from IgD^-^, switched memory resting (IgD-CD21+CD27+) and switched memory active (IgD^-^CD21^-^CD27^+^) subsets were identified(74). Data was acquired on a BD Fortessa flow analyzer, and all data was analyzed using FlowJo v10.10.0.

### Activated-Induced Marker (AIM) Assay

As described in Garrido et. al (75) cryopreserved lymph node cells were thawed, washed, and rested in cRPMI at 37oC with 5% CO2 for three hours. Cells were resuspended in fresh AIM-V Medium (Gibco) at a density of 1×10^6^ cells/mL before being seeded into a 24-well plate at a density of 0.5-2×10^6^ cells per well. Cells were stimulated with BG505 Env peptide pool (1μg/mL) [ARP-13122 Peptide Pool, HIV-1 Subtype A (BG505) Env Protein; 213 peptides, 50ug/ peptide], DMSO (Sigma-Aldrich), or SEB (Staphylococcal enterotoxin B) (0.5μg/mL) for 18-20 hours. Cells were washed, surface-stained (**S4 Table**), fixed with 1% PFA, and acquired within 4 hours on the LSRFortessa running BD FACSDiva Software v.8.0.1 (BD Biosciences) and analyzed using Flowjo Software v.10.10.0.

### T cell Phenotyping

Cryopreserved PBMCs were thawed, washed, and resuspended in cRPMI containing anti-CD28 (0.5 µg/mL) and anti-CD49d (0.5 µg/mL). Cells (1-2×10^6^) were then stimulated for 6 hours with BG505 Env peptide pool (1μg/mL), DMSO (Sigma-Aldrich), or Cell Stimulation Cocktail (0.5x, Invitrogen) at 37°C with 5% CO_2_. Brefeldin A (1x, Invitrogen) was added one hour into the incubation. After stimulation, cells were washed, surface-stained, permeabilized, and then stained for intracellular markers (**S5 Table**), as further described in (75). Stained cells were acquired on the Aurora instrument running SpectroFlo v.3.0.3 (Cytek Biosciences). Flow cytometry was analyzed using Flowjo Software v.10.8.1.

### Binding Antibody Multiplex Assay (BAMA)

HIV-1 antigens, AMC008v9, AMC009v9, AMC011v9, AMC016v9, TRO11v9 were conjugated to magnetic beads (Luminex) as previously described(32) and added to filter plates (Millipore Sigma) with either plasma or monoclonal antibodies as controls (B121R: 1000ug/mL, and Rhesus HIV IgG (Rhivg): 1048.6ug/mL]). Plasma was diluted 1:100, and the monoclonal antibodies were serially diluted in diluent solution (1% powdered milk, 5% goat serum, and 0.05% tween-20 in 1XPBS, pH7.4). Samples were then incubated with the beads for 30 minutes while shaking at room temperature. To detect IgG binding, a PE-conjugated mouse anti-monkey IgG (Southern Biotech) was used at 4ug/mL. The beads were washed and acquired on a Bio-Plex 200 instrument (BioRad) where binding responses were reported at mean fluorescent intensity (MFI). Background was determined by assessing the MFI of binding to blank wells and non-specific binding of samples to a conjugating blank bead during the analysis. An HIV-envelope specific antibody response was considered positive if the samples had an MFI above the lower limit of detection (100 MFI).

### IgG Isolation and Surface Plasmon Resonance (SPR)

IgG was isolated from plasma using Protein G plates (Cytiva). Plasma samples were diluted 1:1 with 1x TBS and filtered through Spin-X Centrifuge Tube filters (Costar) and then added to the Protein G plates. Samples were incubated on the plate for at least an hour while shaking at room temperature. After the IgG-depleted plasma was spun out, the IgG was extracted from the plates by adding 2.5% acetic acid and eluting them into 1M Tris-HCl pH 8.0. The purified IgG was passed through 30K Centrifugal Filters (Amicon), and the buffer was exchanged with 1X PBS.

Avidity of purified IgG against BG505 GT1.1 SOSIP-his trimer was measured by SPR on a BIAcore T200 instrument (Cytiva) at 25°C (24,76). HBS-EP^+^ (0.01M HEPES pH7.4, 0.15M NaCl, 3mM EDTA, 0.05% v/v Surfactant P20) was used as a running buffer throughout. Briefly, A series S CM3 sensor chip was covalently conjugated with anti-His antibody by amine coupling chemistry His-tagged, Env trimer was immobilized on the sensor surface to R_L_ values giving R_max_ values close to 15 response units (RU). Trimer was captured in two flow cells (Fc2 and Fc4), while two flow cells (Fc1 and Fc3) served as reference. Purified IgG samples were run with single-cycle kinetics (SCK). Five concentrations were injected in ascending order up to 250 nM by two-fold diluted samples. Association was monitored for a cumulative 300(s) followed by a dissociation phase of 3600(s) at the end of the highest concentration. The surface was regenerated by injecting glycine (10nM; pH 1.5) 3 times with a contact time of 90(s). As a control, recombinant bnAb, VRC01 was injected at two-fold ascending concentrations up to 50nM. Throughout the binding experiments, a high flow rate (50µl/min) was used to avoid mass transport limitation. In each batch of experiments, 2-3 start-up cycles and 4-5 cycles of buffer injections (zero analyte) were included.

Non specific binding of IgG to reference flow cell was checked prior to actual binding experiment. Binding data generated was double-reference subtracted (reference flow cell and buffer injection with zero analyzte). Data analyses were done with BIA evaluation software [v3.2.1] (Cytiva). A Langmuir model was applied to determine kinetic parameter and fitted satisfactorily.

Binding responses were measured by average post-injection response unit (RU) over a 10s window at the top of the association phase, and the dissociation rate constant, k_off_(s^-1^), was measured during the post-injection phase. Positive response was defined when both replicates had a RU value ≥ 15. Relative avidity binding score is calculated as follows: Avidity score (RU.s) = (Binding RU/k_off_(s^-1^))(24,77,78).

### Fc Glycan Analysis

IgG Fc-N-linked glycan analysis was performed as previously reported (79). Briefly, EZ-Link NHS-LC-LC-Biotin (Thermo Fisher Scientific, #21323) was used at a 20:1 biotin-to-protein molar ratio to biotinylate BG505 GT1.1. Biotinylated BG505 GT1.1 SOSIP were incubated with streptavidin magnetic beads (NEB, #S1420S), which were later washed twice with washing buffer (0.5 M NaCl, 20 mM tris-HCl, and 1 mM EDTA; pH 7.5) to remove the unbound BG505 GT1.1 in the supernatant.100 μl of antigen-free magnetic beads were incubated with a total of 200 μl of rhesus macaque serum to pull down nonspecific binding serum glycoproteins. Sera was then incubated in the precleaned serum with 200 μl of the BG505 GT1.1 -coupled beads at 4°C overnight to allow the binding of antigen-specific antibodies. Rhesus IgG Fc region was released by incubating BG505 GT1.1 -specific antibody-bound beads with IdeZ (NEB, #P0770S) at 37°C for one hour. An excess amount of protein A on Protein A HP MultiTrap (Cytiva, #28903133) was used to purify total serum IgG. The Fc region was separated from Fab with Protein A HP MultiTrap after digestion with IgG protease. The procedure of deglycosylation and glycan labeling was performed following the manufacturer’s instructions (Agilent Technologies Inc., #GX96-IQ). The removal of tris-HCl in the IgG protease buffer was conducted via Buffer exchange to 20 mM Hepes. The labeled glycans were analyzed using the Gly-Q Glycan Analysis System GQ2100 (ProZyme now Agilent Technologies Inc.). The built-in software automatically generated the relative abundance of each glycan species to minimize human error in the data processing. To calculate the relative abundance of each glycan species, the peak area of the given glycan was divided by the areas of all the peaks on the electropherogram summed together: relative abundance = (peak area of a glycan/area summation of all peaks observed) × 100. A subset of samples was manually set to the detection cut-off (0.05% peak area) due to the absence of the relative abundance of some glycan species.

### Statistical analysis

To compare IgG plasma binding between the infants and juveniles, trapezoidal AUC was calculated using ED50 values from ELISA from week 0 to week 150. AUC between two subsequent time points was computed by averaging ED50 values, subtracting the lower limit of detection (LLOD), and multiplying by the time interval. All trapezoidal AUCs were summed to obtain total AUC for each subject. The total AUC was, then, divided by the number of weeks per subject. For statistical comparisons, two-sided unequal variance t-test (Welch’s t-test) was used to compare total AUC and ID50 values from neutralization data after log-10 transformation. For the comparison of frequency of animals with greater than three fold-change in ID50 titers, the Fisher’s exact test was performed. In the secondary endpoint and exploratory analysis, Avidity, BAMA, and ADCP data were first transformed by log-10, log-10, and log-e, respectively based on its original scale, then two-sided Welch’s t-test was performed. Proportions from TFH/AIM, Ag-specific B cells, and Glycan data were compared using two-sided Wilcoxon-rank sum tests with normal approximation. Unadjusted P values are reported for primary and secondary endpoints. In exploratory analysis, Benjamini-Hochberg adjusted P values to account for multiple comparisons was reported within each data type as well as unadjusted P values.

## Acknowledgments

We would like to thank J. Watanabe, J. Usachenko, and the staff of the CNPRC Colony Research Services and Clinical Laboratories for the support with these studies. Flow cytometry was performed in the WCM Flow Cytometry Core Facility (New York, NY) or the UNC Flow Cytometry Core Facility (Durham, NC). We thank T. Bijl and J. Snitselaar of Amsterdam UMC, Academic Medical Center, for the production and purification of the BG505 SOSIP trimers. We are also grateful for the PAVEG contract no. HHSN272201800004C for supporting this work. We thank Leonidas Stamatatos and Andrew T. McGuire for the 426c pseudoviruses. The 3M-052-SE was provided by AAHI and 3M. The funders had no role in the study design, data collection, interpretation, or the decision to submit the work for publication. The content is solely the responsibility of the authors and does not necessarily represent the official views of the NIH. Study data were collected and managed using REDCap (Research Electronic Data Capture) electronic data capture tools hosted at Duke University and Weill Cornell Medicine. The figure of the study design was created with BioRender.com.

## Contributions

S.R.P., K.D.P., K.K.A.V.R., K.W., M.G.H., J.P.M., R.W.S., D.C.M., R.D., and A.N.N. contributed to the conception and study design. Y.A.I, X.S, J.E., M.D, and J.I performed ELISA assays. Y.A.I performed plasma BAMA assays. Y.A.I and A.Y performed SPR on isolated IgG from plasma samples. K.V. performed ADCP assays. X.S and D.M. performed sera neutralization screening. L.M.S. and S.Z performed IgG EMPEM assays. ADCC was performed under S.S.O and G.F. D.D. performed T cell phenotyping and AIM assays. Y.A.I performed B cell phenotyping assays. J.C. performed IgG antibody Fc glycosylation assays. E.K. and M.H. performed statistical analyses. Y.A.I., X.S., G.O., E.K., M.G.H., P.J.K., J.P.M., R.W.S., S.R.P., K.D.P., and A.N.N. contributed to the interpretation of the results. J.E., C.W., H.G., S.M.A., A.B.W., D.C.M., A.A.N., S.R.P., K.D.P., K.K.A.V.R. were involved in supervision or project administration. Y.A.I., S.R.P, and A.N.N were responsible for drafting the original manuscript. All authors were involved in critical revision of the manuscript for accuracy and important intellectual content.

---

**Supp Fig 1.**
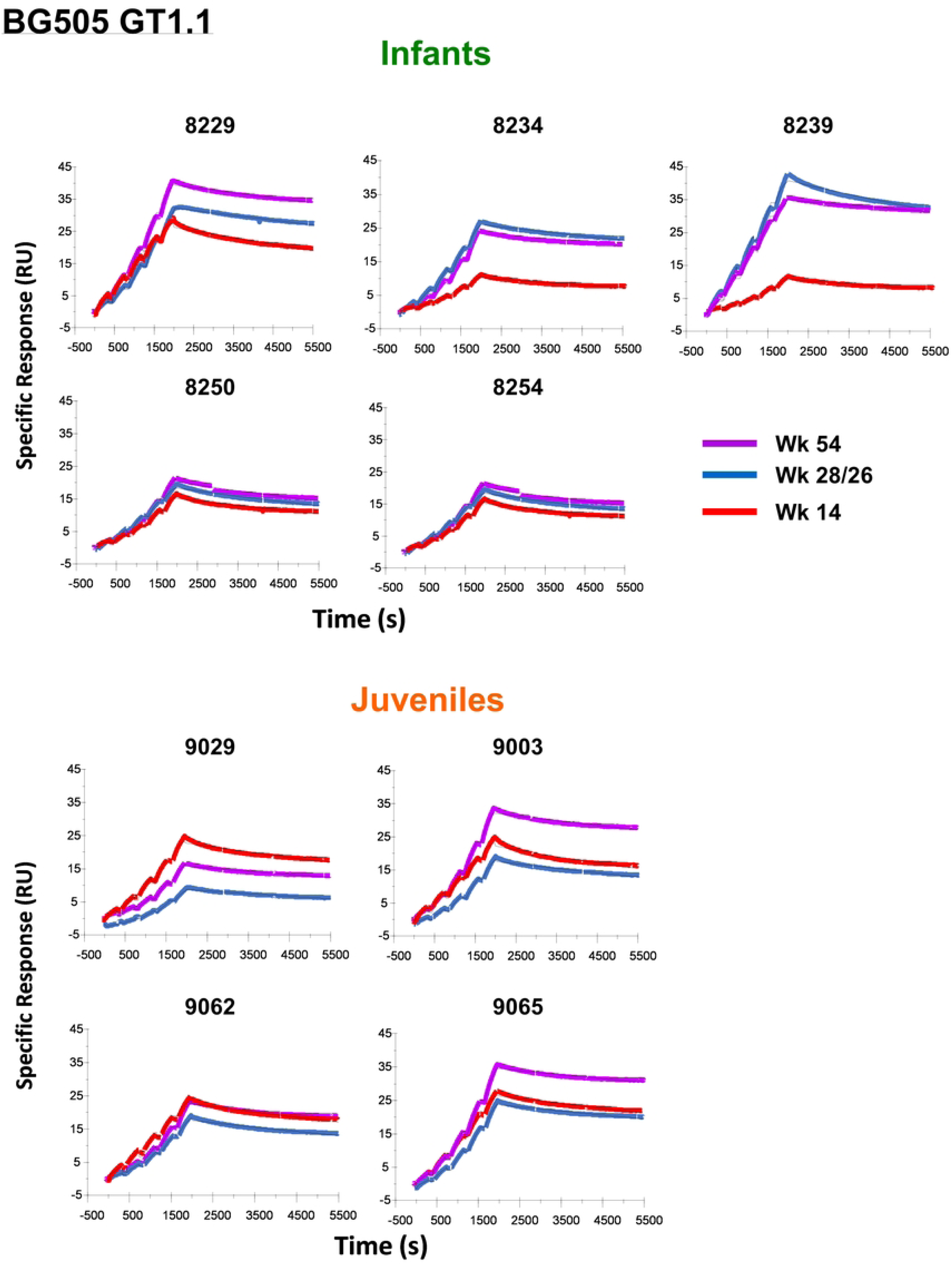

**Supp Fig 2.**
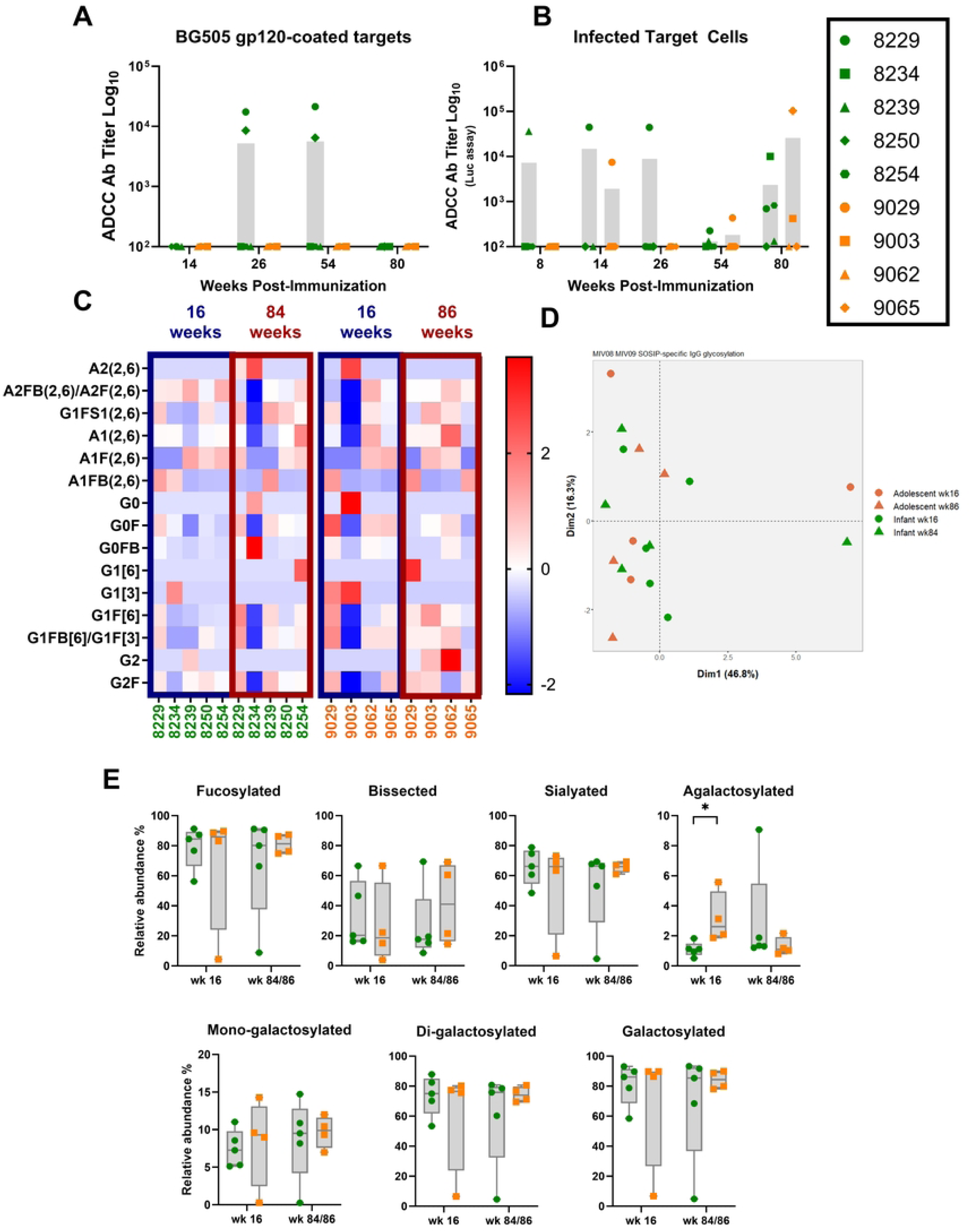

**Supp Fig 3.**
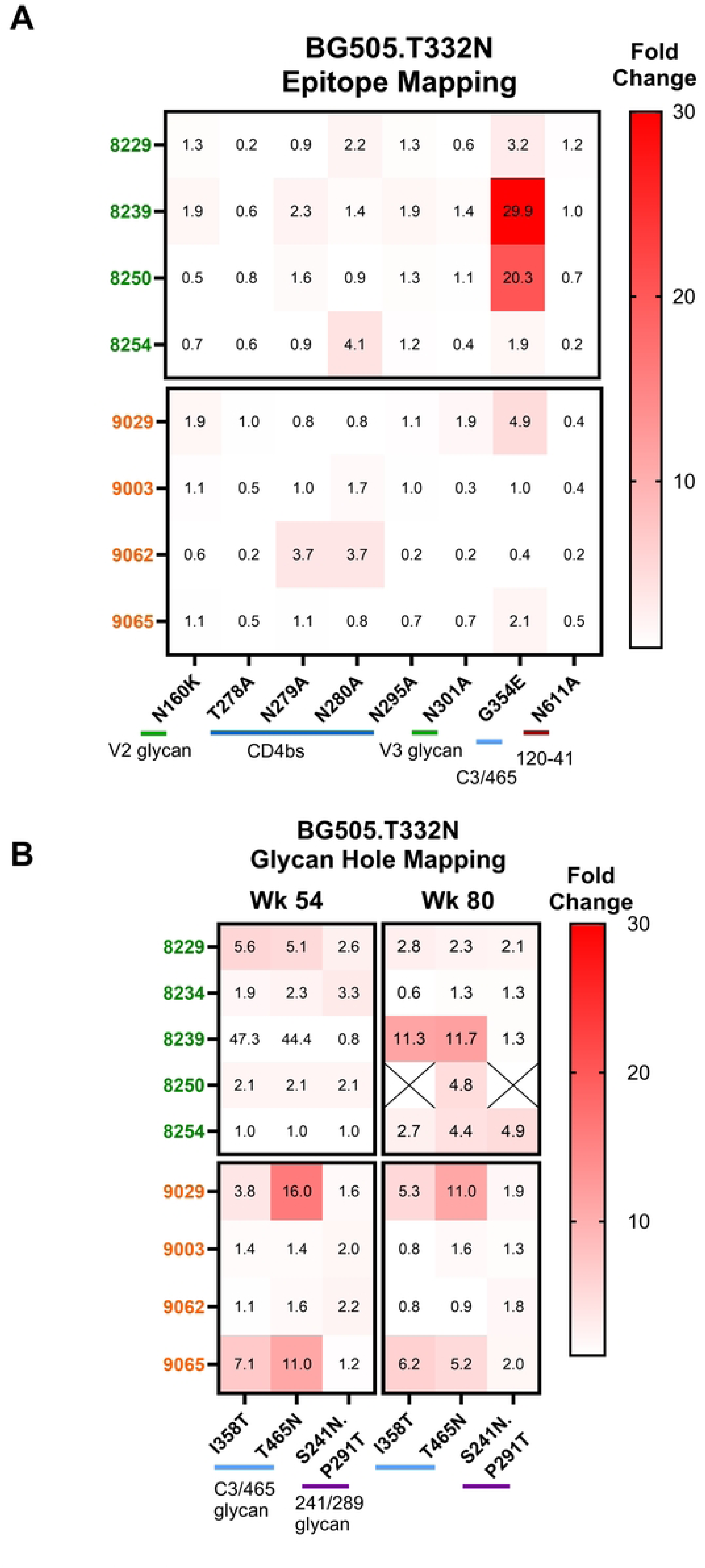

**Supp Fig 4.**
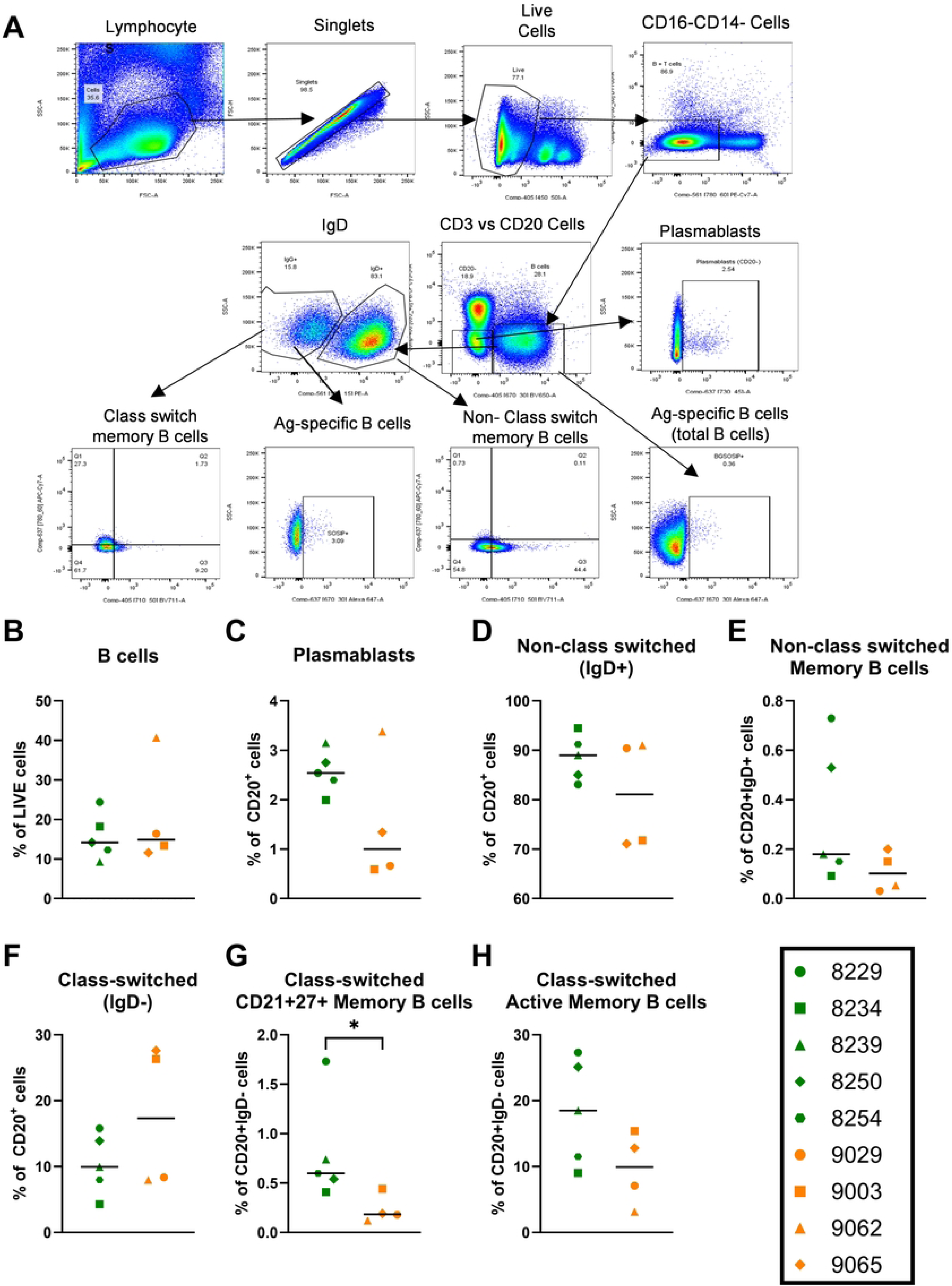

**Supp Fig 5.**
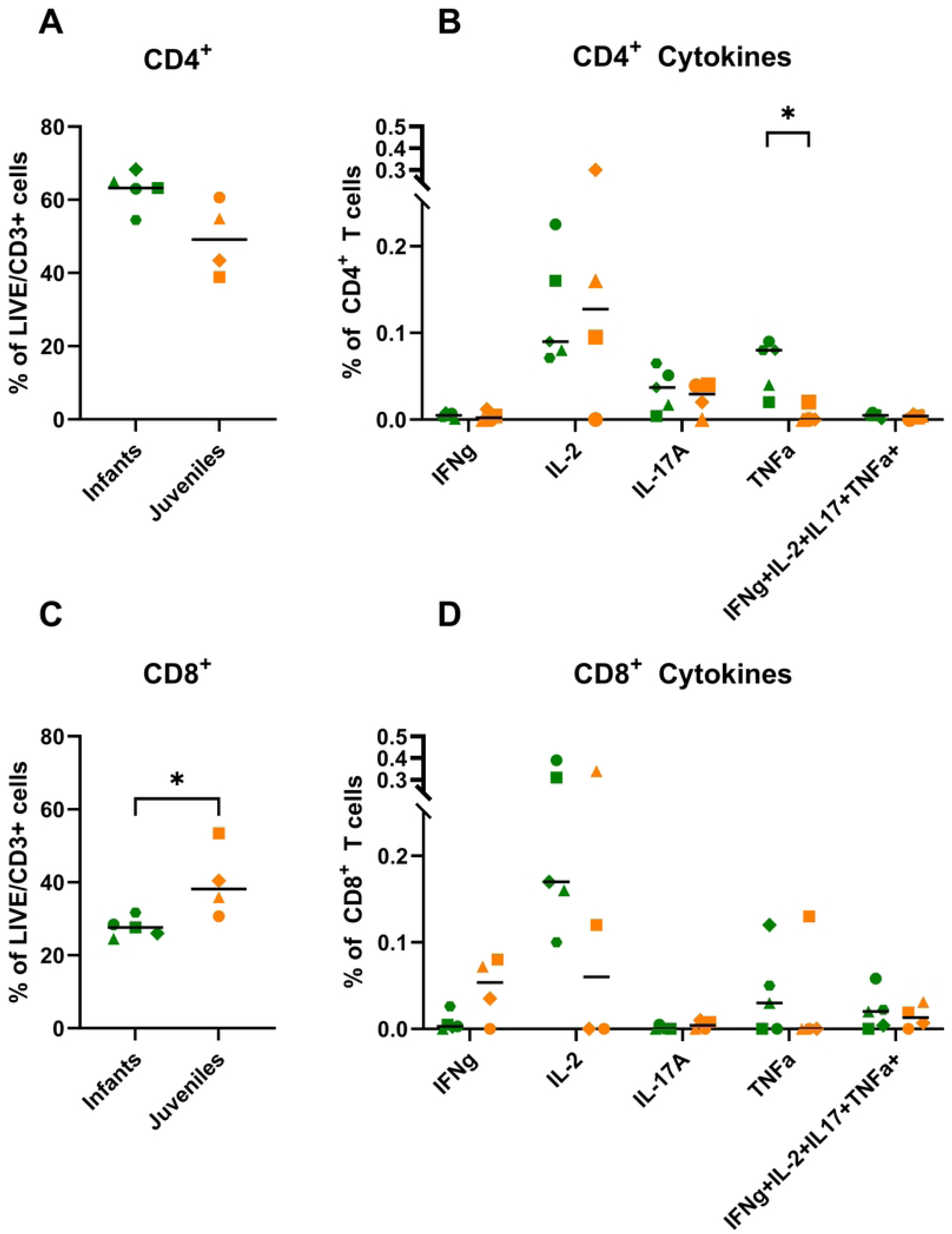

**Supp Fig 6.**
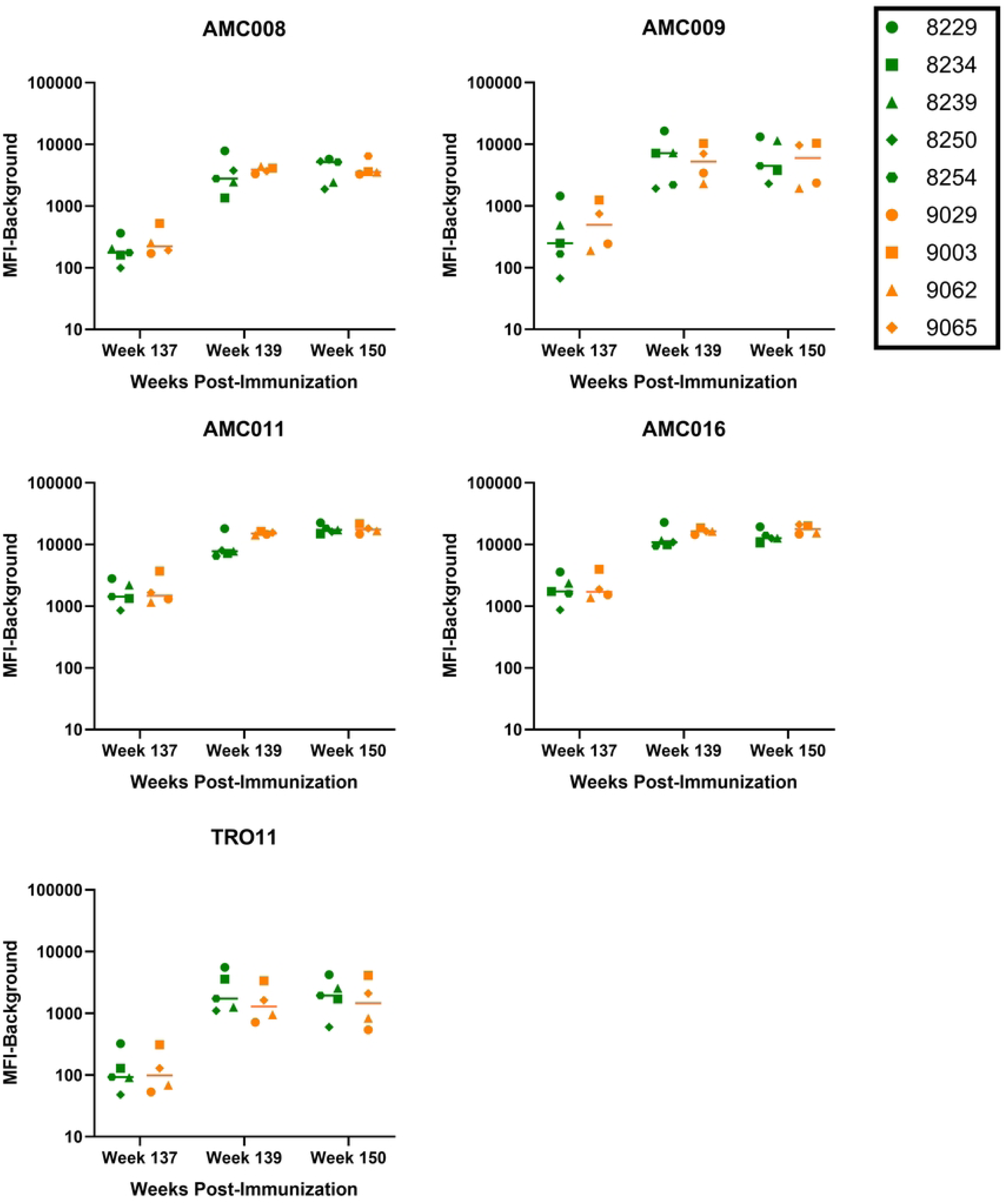

